# Revisiting atypical language lateralization in dyslexia

**DOI:** 10.1101/2025.05.06.652381

**Authors:** Helena Verhelst, Emma M. Karlsson, Robin Gerrits, Guy Vingerhoets

## Abstract

Hemispheric lateralization has been central to developmental dyslexia research for over a century, yet its role in the etiology of reading and language deficits remains elusive. While altered asymmetries have long been implicated, evidence is inconsistent, with limited consideration given to individual variability in lateralization patterns.

This study investigated hemispheric lateralization in 35 adults with dyslexia and 35 matched controls using functional MRI across three language tasks: word generation, rhyming decision, and lexical decision. Laterality indices (LIs) were calculated to comprehensively assess the strength, direction, and consistency of activation across global and regional task-specific brain areas.

Significant group differences were not found in the absolute strength of lateralization for global measures or any regional measures, except in the fusiform gyrus, where people with dyslexia showed lower asymmetry. Directional asymmetry was similar across the two groups, except in the fusiform gyrus during the reading task, where dyslexic individuals showed a higher prevalence of right-hemisphere lateralization compared to controls. Interestingly, we found that dyslexic participants demonstrated greater inconsistency in regional lateralization during reading and rhyming tasks. Among individuals with dyslexia, those with inconsistent lateralization in the reading task had weaker fusiform lateralization, although fusiform LI strength itself did not predict reading outcomes.

Our findings suggest that dyslexia is characterized by inconsistent, rather than universally weaker, lateralization patterns. Inconsistencies in task-related and regional lateralization may disrupt the efficiency of language networks, contributing to observed reading deficits. By highlighting the role of regional and task-specific inconsistencies, this study provides new insights into the neural mechanisms underlying dyslexia and underscores the importance of considering individual variability in hemispheric lateralization when investigating language disorders.

## Introduction

Developmental dyslexia, commonly referred to as dyslexia, is a neurodevelopmental learning disorder defined by persistent reading and spelling difficulties, despite otherwise normal development.^1,2^ A core characteristic of dyslexia is a deficit in phonological decoding, which disrupts the representation, storage, and retrieval of speech sounds^3^. These difficulties are commonly identified during school age but typically persist into adulthood.^4,5^ Although some symptoms may evolve over time and individuals frequently develop coping strategies, core difficulties with reading, spelling, and language processing generally remain,^6^ significantly impacting quality of life.^7–9^

Given the persistent nature of dyslexia and its profound impact on daily functioning, considerable effort has been devoted to understanding its neurobiological basis. One key area of focus has been hemispheric lateralization, particularly given the left hemisphere’s well-established dominant role in language processing.^10,11^ Early theories of dyslexia proposed that atypical hemispheric lateralization for language might play a fundamental role in its development (e.g., ^12–14)^. Notably, Orton’s 1925 seminal theory^13^ suggested that dyslexia symptoms result from a failure to establish hemispheric lateralization for language. However, early behavioral studies employing techniques such as visual half-field and dichotic listening to examine lateralization in dyslexia produced inconsistent findings. Some studies reported no differences in language asymmetry,^15–17^ while others found evidence of reduced or even reversed lateralization in dyslexia groups.^18,19^ Of course, these behavioral measures provide only indirect evidence of lateralization, relying on performance differences rather than directly assessing neural activity.

The advent of functional neuroimaging techniques has enabled more direct investigations into the neurobiology of dyslexia, including hemispheric asymmetry, by inferring brain activity from hemodynamics. Reviews of whole-brain functional MRI studies have identified altered activation in several brain regions associated with reading and phonological processing, including the left occipito-temporal cortex, left temporo-parietal cortex, and left inferior frontal gyrus, in both children and adults with dyslexia (e.g., ^20–26^). However, despite this evidence suggesting hypoactivation in left hemisphere regions, surprisingly few neuroimaging studies have *specifically* focused their investigations on functional lateralization in individuals with dyslexia. Conventional activation map comparisons, which are performed on threshold-dependent group averages, are insufficient for examining hemispheric lateralization as asymmetrical differences vary significantly with threshold changes.^27–29^ Instead, researchers should explicitly examine hemispheric contributions within individuals, which is typically performed using a laterality index (LI).^27^ The LI quantifies both the degree (strength) and direction (left vs. right) of hemispheric dominance and is calculated by subtracting activation levels between the left and right hemispheres within an individual, either across the whole brain or within specific regions of interest.^30^

Even in the very few studies where asymmetries are quantified in individuals with dyslexia, findings are inconsistent. For instance, Hernandez et al.^31^ used fMRI to compare 15 adults with dyslexia to 15 controls on a phonological rhyme judgment task, measuring asymmetry in six regions of interest. While the control group had left-hemisphere LIs in all regions, the dyslexia group did not lateralize in two of these regions. The rest of the regions had asymmetries comparable to controls. Based on these findings, the authors concluded that individuals with dyslexia lack asymmetry in certain language regions. Similarly, Waldie et al.^32^ reported a lack of left-hemisphere lateralization in the temporal lobe during lexical decision tasks in a group of 12 adults with dyslexia compared to 16 controls. However, unlike Hernandez et al.^31^ they observed typical left lateralization in the frontal lobes.

A key limitation of these studies, is the reliance on group-averaged LI values as it conceals important individual differences.^33–35^. LI values typically range from negative to positive, reflecting both the direction (left versus right) and degree (weak versus strong) of asymmetry, but this scale must be carefully interpreted when drawing conclusions from group averaged data. For example, a reduced LI in the dyslexia group may indicate a general decrease in asymmetry across all individuals, or it could be driven by a subgroup having reversed, right-hemisphere lateralization.^36^ Making this distinction in the sample is crucial to fully understand the mechanisms underpinning altered asymmetry in dyslexia. One study that accounted for the possibility of a higher prevalence of right-hemisphere dominance in dyslexics confirmed less lateralized activation in the dyslexia group,^37^ but was unable to provide more regional specificity due to the use of functional transcranial Doppler ultrasonography (fTCD), which only allows for an overall lateralization measure for the left and right middle cerebral arteries. To summarize, while Hernandez et al.^31^ and Waldie et al.^32^ used fMRI to identify regional differences in lateralization, they disagreed on which regions were affected and did not account for individual differences. Whereas Illingworth and Bishop^37^ controlled for individual differences, their method did not allow the evaluation of regional specificity.

To add an extra layer of complexity, it may also be that various task types may result in distinct lateralization profiles in the dyslexia group, potentially contributing to the heterogeneity and lack of consensus on asymmetry in the literature. Bishop et al.^38^ suggested that the lateralization of each language function is seen as an independent probabilistic process, which means some functions may follow the typical leftward bias, while others do not. Increased variability could lead to inconsistency in lateralization across different language tasks, resulting in a less efficient language network, as it requires greater hemispheric integration and communication across the corpus callosum. From this perspective, it may not only be the degree of lateralization, as early theories suggested, but also (in)consistency of lateralization across the two hemispheres that give rise to symptoms seen in dyslexia. Some evidence supporting this idea comes from an fTCD study on individuals with developmental language disorders, including individuals with dyslexia. In this study, Bradshaw et al.^39^ did not find any group differences in LI scores between the developmental language disorder (DLD) group and control group, as found by Illingworth and Bishop.^37^ However, they did observe that individuals with DLD displayed greater inconsistency in side of lateralization (left or right) across their three language tasks (phonological decision, semantic decision and sentence generation) compared to individuals in a control group.

Here, we revisit the century-old question of the role of lateralization of language functioning in dyslexia, by carefully considering individual differences in our sample. More specifically, we used fMRI to measure hemispheric lateralization across three tasks - word generation, rhyme judgment, and lexical decision - each chosen to capture different aspects of language processing. Our study is the first to systematically investigate both the consistency of lateralization across tasks and within task-specific regions of interest (ROIs) in individuals with dyslexia. Furthermore, unlike most previous studies, we distinguish between the strength and direction of lateralization. We predict that, for each of the three language tasks, individuals with dyslexia will (1) be less lateralized, both globally and regionally, (2) show a higher prevalence of right hemisphere dominance, both globally and regionally, and (3) predict that individuals with dyslexia will show more inconsistency in their lateralization, both across different language tasks as well as across different regions within a single language task. In addition, we explore whether alterations in hemispheric asymmetry are associated with reading performance.

## Materials and methods

### Participants

Thirty-five adults with a diagnosis of dyslexia and thirty-five individuals without any diagnosis of language or reading impairment were recruited for this study via social media, word of mouth, and a student participation platform. Inclusion criteria were Dutch as a first language, age between 18 and 40 years, and right-handedness. All participants had normal or corrected-to-normal vision and reported no history of brain injury or neurological disease. All participants in the dyslexia group had been diagnosed by a trained professional, either during their school years or in the context of receiving disability services at university or college. The groups did not differ significantly in terms of age, sex distribution, or years of formal education (all *P* > 0.05, see Table 1). The sample size of 35 adults with dyslexia and 35 adults without was chosen based on feasibility and alignment with similar studies investigating functional asymmetry in neuropsychological populations. While no formal power analysis was conducted, this sample size was considered sufficient to detect meaningful differences in language functioning between groups and is consistent with sample sizes used in comparable research in this field. The present study was carried out in accordance with the ethical rules for human subjects as described in the Declaration of Helsinki. The study was approved by the medical ethics committee of Ghent University Hospital (approval number BC-09822). Written informed consent was obtained from each participant.

**Table 1.** Demographic and behavioural test performance summary statistics for the control and dyslexia groups.

|  | CON |  | DYS |  | *P*-value | Effect size |
| --- | --- | --- | --- | --- | --- | --- |
|  | *M* | *SD* | *M* | *SD* |  |  |
| Sex (male/female) | 13/22 |  | 14/21 |  | 0.843 | -0.16 |
| Age (years) | 23.26 | 5.08 | 23.40 | 5.33 | 0.909 | -0.03 |
| Education (years) | 14.60 | 1.91 | 13.83 | 2.01 | 0.104 | -0.39 |
| Word reading test | 109.40 | 16.82 | 72.14 | 16.07 | **<0.001** | **2.27** |
| Spelling test<sup>a</sup> | 125.30 | 8.97 | 95.97 | 13.96 | **<0.001** | **2.49** |
| Spoonerism test<sup>b</sup> | 52.12 | 16.14 | 108.61 | 48.46 | **<0.001** | **-1.56** |

CON = control group DYS = dyslexia group, *M* = Mean, *SD* = standard deviation, *P*-values are for comparisons between the control and dyslexia groups (two-proportions *z* test for sex, two-sided independent *t*-test for the other demographics), effect sizes are Cohen's *h* for sex and Cohen's *d* for the other demographics and tests.
<sup>a</sup> Values for the spelling test for the control group are shown for *n* = 34 as one control participant did not complete this test
<sup>b</sup> Spoonerisms: lower score equals a better performance

### Behavioural measures

Participants completed three language tests that have shown to suffice to correctly diagnose dyslexia in adults: word reading, word spelling, and phonological awareness.^40^ In the Word Reading Test for Students,^41^ participants were required to read aloud as many words as possible within one minute. The word spelling subtest of the GL&SCHR involved participants writing down 30 exception words that were read out to them.^42^ For this test, a weighted score that accounted for both accuracy and confidence in spelling was computed. Phonological awareness was evaluated using the Spoonerisms subtest from the GL&SCHR,^42^ where participants had to switch the first letters of two auditorily presented words, such as transforming ‘Harry Potter’ into ‘Parry Hotter’. Here, again, a weighted score factored in both accuracy and reaction time.

### MRI paradigms

#### Lexical decision task

Word reading was assessed using a lexical decision task in a block design (hereafter referred to as ‘reading’). During the experimental blocks, participants were instructed to press a button with both their left and right index fingers simultaneously if the stimulus was a word, and to refrain from pressing if the stimulus was a non-existent word. During the control blocks, participants were tasked with determining whether lines displayed on the screen were all oriented in the same direction. If they were, participants pressed a button with both index fingers; if not, they withheld the button press. Stimuli in the experimental blocks consisted of 24 words and 24 non-words. Words were 4–7 letters long, with six words of each word length. The pseudowords matched the words in length and were created using Wuggy.^43^ Control block stimuli consisted of strings of slashes and backslashes matching the length of the (pseudo)words in the experimental conditions (e.g., all in the same direction: ‘////’, not all in the same direction: ‘//\///’). All stimuli were displayed once in a random order in the font ‘Courier New’, in black on a white background (see Figure 1 for an example trial). Each trial started with a fixation cross shown in the centre of the screen. After 500 ms, a stimulus appeared for 800 ms, followed by a dash that remained visible for 1 s. The experiment consisted of six experimental and six control blocks, each containing eight trials. An instruction screen appeared for 2 s at the beginning of each block with a one-word prompt (‘woord?’ [word?] or ‘zelfde?’ [same?] for the experimental and control block, respectively). These blocks alternated with 15 s rest periods, during which a dash was displayed.

**Figure 1.**
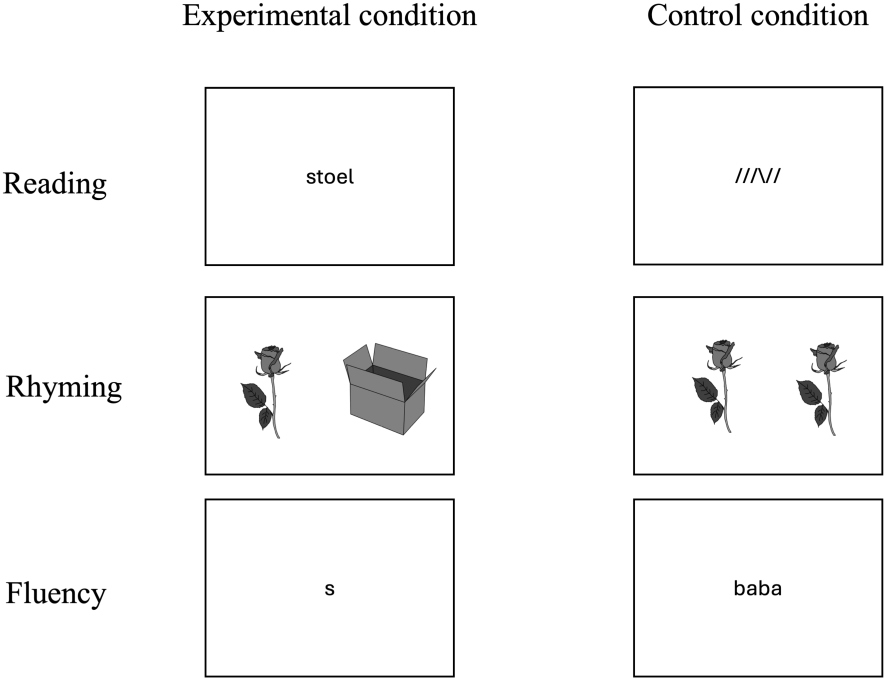
An example trial from the experimental and control conditions for each of the three functional MRI paradigms.

#### Rhyming task

Phonological awareness was examined using a rhyming task in a blocked design (hereafter referred to as ‘rhyming’). During the experimental blocks, participants were instructed to press a button with both index fingers simultaneously if the stimulus pair formed a rhyme and to refrain from pressing if the pair did not form a rhyme. During the control blocks, participants were asked whether the stimulus pair was an exact match. If the answer was yes, they had to press a button with both index fingers; if not, they withheld the button press. The stimuli consisted of 84 black-and-white line drawings, each depicting highly familiar, high-frequency monosyllabic words in spoken Dutch. The line drawings were taken from the Dutch-Belgian version of the Multipic pictures and names.^44^ Forty-two pictures were combined to form 21 rhyming pairs. Non-rhyming trials were created by pairing the remaining 42 items such as no phonological overlap existed (e.g., ‘koe’ – ‘wolk’). The stimuli used in the rhyme blocks were also used in the ‘same picture?’ control task. Twenty-one of the pictures were presented as identical pairs, while another 42 pictures were re-paired to form ‘different picture?’ trials. Thus, of the 84 pictures seen in the rhyme condition, 63 were also presented in the ‘same picture?’ control condition. During each trial, a picture pair was shown on the screen for 3 s, during which participants could respond (see Figure 1 for an example trial). The experiment consisted of seven experimental and seven control blocks, each containing six trials. An instructions screen appeared for 2 s at the beginning of each block with a one-word prompt (‘Rijm?’ [rhyme?] or ‘Zelfde?’ [same?] for the experimental and control block, respectively). These blocks alternated with 15 s rest periods, during which a dash was displayed.

#### Verbal fluency task

Lexical retrieval was assessed using a covert phonemic verbal fluency task (hereafter referred to as ‘fluency’) in a blocked design. During the experimental blocks, participants were asked to think of as many words as possible that began with a letter displayed in the centre of the screen. The seven letters b, d, k, m, p, r, and s were used. In the control blocks, participants silently repeated the meaningless string ‘baba’ shown on the screen (see Figure 1 for an example trial). The experiment consisted of six experimental and six control blocks, each with a duration of 15 s. These blocks alternated with 15 s rest periods, during which a dash was displayed in the centre of the screen.

### MRI Acquisition

The scans were acquired in a Siemens 3 Tesla Prisma magnetic resonance (MR) scanner located at Ghent University Hospital, using a 64-channel head coil. All functional images were acquired with a T2-weighted gradient-echo EPI sequence, multiband factor: 4, field of view (FOV; mm) = 210 x 210, 60 slices; acquired voxel size (mm) = 2.5 x 2.5 x 2.5 repetition time (TR) = 1070 ms, echo time (TE) = 31 ms, flip angle (FA) = 52°. The number of acquisition volumes for each task were as follows: 393 for the word generation task, 458 for the rhyme task, and 397 for the reading task. The field map was acquired with the following parameters: FOV (mm) = 210 x 210, 60 slices; acquired voxel size (mm) = 2.5 x 2.5 x 2.5, TR = 588 ms, TE = 4.92 ms and 7.38 ms, FA = 60°. T1-weighted structural images were obtained using a MPRAGE sequence with the following scan parameters: TR = 2250 ms, TE = 4.18 ms, inversion time [TI], 900 ms, FA = 9°, FOV (mm) = 256 x 256, 176 slices, voxel size (mm) = 1 x 1 x 1.

### MRI processing

#### fMRI preprocessing

Preprocessing was performed using *fMRIPrep* 23.2.1^45^, which is based on *Nipype* 1.8.6^46^.

##### Preprocessing of B0 inhomogeneity mappings

A *B0* nonuniformity map (or *fieldmap*) was estimated from the phase-drift map measure with two consecutive gradient-recalled echo acquisitions. The corresponding phase-map was phase-unwrapped with prelude (FSL None).

##### Anatomical data preprocessing

The T1-weighted (T1w) image was corrected for intensity non-uniformity (INU) with N4BiasFieldCorrection^47^, distributed with ANTs 2.5.0^48^, and used as T1w-reference. The T1w-reference was skull-stripped with a *Nipype* implementation of the antsBrainExtraction.sh workflow (from ANTs), using OASIS30ANTs as target template. Brain tissue segmentation of cerebrospinal fluid (CSF), white-matter (WM) and grey-matter (GM) was performed on the brain-extracted T1w using fast^49^ (FSL). Volume-based spatial normalization to standard space (MNI) was performed through nonlinear registration with antsRegistration (ANTs 2.5.0), using brain-extracted versions of both T1w reference and the T1w template. The *ICBM 152 Nonlinear Asymmetrical template version 2009c*^50^ was selected for spatial normalization and accessed with *TemplateFlow*^51^ (23.1.0).

##### Functional data preprocessing

For each of the three BOLD runs, the following preprocessing was performed. First, a reference volume was generated, using a custom methodology of *fMRIPrep*, for use in head motion correction. Head-motion parameters with respect to the BOLD reference (transformation matrices, and six corresponding rotation and translation parameters) were estimated before spatiotemporal filtering using mcflirt (FSL, Jenkinson et al. 2002). The estimated *fieldmap* was then aligned with rigid-registration to the target EPI reference run. The field coefficients were mapped on to the reference EPI using the transform. The BOLD reference was then co-registered to the T1w reference using mri_coreg (FreeSurfer) followed by flirt (FSL, Jenkinson and Smith 2001) with the boundary-based registration (Greve and Fischl 2009) cost-function. Co-registration was configured with six degrees of freedom. Gridded (volumetric) resamplings were performed using nitransforms, configured with cubic B-spline interpolation. After preprocessing in *fMRIPrep*, analysis was continued in SPM12 running in MATLAB_R2019b. Here, normalized data was spatially smoothed with a 5mm^3^ full-width half maximum Gaussian kernel.

#### fMRI analysis

In SPM12, the general linear model was used to predict the hemodynamic response curve by applying regressors that coded for each experimental and control condition, along with the six movement parameters (three rotation and three translation) estimated during preprocessing. A boxcar function, convolved with the hemodynamic response function, was fitted to the time series at each voxel, resulting in weighted beta images. The beta images of the experimental and control condition were then contrasted to create a t-statistic map.

##### Determination of Participant-Specific LIs

The LI-toolbox plugin^29,52^ for SPM was used to assess hemispheric contribution based on the t-statistic map for each of the three tasks. This toolbox allows for comparisons between the right and left hemispheres without commonly cited problems such as complications that arise from statistical outliers, threshold-dependent comparisons, or data sparsity.^52^ The toolbox employs a bootstrapping method to calculate a LI ranging from -1 to +1 using the standard LI formula (LI = (L-R)/(L+R)). Note that we inverted the scores to be in accordance with the Laterality indices consensus initiative (LICI),^30^ thus, a negative score indicates greater *left hemisphere* activation, and a positive score indicates greater *right hemisphere* activation. Participant-specific LIs were calculated for task-specific regions of interest (ROIs; see below) for each of the three fMRI tasks, using the recommended settings.^52^

##### Regions of interest

The ROIs were selected from the Automated Anatomical Labelling (AAL) atlas.^53^ We included regions known to be lateralized for the examined function based on previous research. For the reading task, we selected the inferior frontal opercularis + triangularis (Broca’s area, hereafter referred to as ‘inferior frontal gyrus’), precentral gyrus, middle temporal gyrus, fusiform gyrus, inferior parietal gyrus, and inferior occipital gyrus.^54^ For the rhyming task, the inferior frontal gyrus, precentral gyrus, middle temporal gyrus, supplementary motor area, and cerebellum were selected.^55–61^ Finally, for fluency we chose the inferior frontal gyrus, precentral gyrus, supplementary motor area, thalamus, and cerebellum.^62^ The ROIs were picked from the AAL atlas using the WFU PickAtlas v 3.0.5^63,64^ and made symmetrical by selecting the left and right ROI, flipping them over the x-axis and then adding them up. To create the global ROIs, we summed the task specific ROIs for each task separately.

### Statistical analyses

To examine group differences in **strength** of lateralization, we used t-tests on the absolute LI values. Given the hypothesis that language lateralization would be decreased in people with dyslexia, all t-tests were one-tailed. FDR corrections were used to correct for the multiple comparisons. Cohen’s d was calculated as a measure of effect size for all tests. The alpha-level that was used to determine significance for all tests was *P* < 0.05.

To examine group differences in the **direction** of lateralization, we used Fisher’s exact tests. With these tests, an association between group (dyslexia versus control) and direction (left versus right) of lateralization was tested. To do this, participants were first classified as either left or right lateralized based on individual LI values (with a negative value categorized as left hemispheric and a positive value as right hemispheric). All tests were one-tailed, as we hypothesized that the dyslexia group would exhibit increased incidence of right hemisphere lateralization compared to controls.

Finally, the group difference in the **consistency** of lateralization was tested using Fisher’s exact tests. Here, the association between group (dyslexia versus control) and consistency (consistent = all tasks/ROIs lateralize to the individual’s ‘typical’ (language dominant) hemisphere versus inconsistent = at least one task/ROI lateralizes to the individual’s ‘atypical’ (language non-dominant) hemisphere) was tested. As we predicted that the individuals with dyslexia would show more inconsistency in their lateralization, a one-tailed approach was also taken here. A dichotomous subdivision (consistent versus inconsistent) is quite coarse when looking at the multiple task-specific ROIs as it does not allow us to see whether the inconsistency is low (i.e. only one ROI is lateralized to the atypical hemisphere) or high (i.e. more than one ROI is lateralized to the atypical hemisphere). That is why in a next step, we further subdivided the inconsistent individuals according to the number of atypical ROIs. Again, Fisher’s exact tests were used to determine if there was an association between group (dyslexia versus control) and degree of inconsistency (one versus two versus three atypical ROIs). It is important to note that for this analysis, the typical hemisphere is defined as the hemisphere where the majority of tasks/ROIs lateralize to. For example, an individual with 2 out of three tasks lateralized to the right hemisphere is defined as a right lateralized individual.

Lastly, to examine if inconsistent lateralization was associated with poorer reading performance, we divided dyslexia participants in two groups: consistent and inconsistent for the reading task and used a t-test to compare the performance on the word reading test. This test was also one-tailed, as we hypothesized worse reading performance in inconsistently lateralized participants.

### Data availability

The MRI data that support the findings of this study are openly available in OpenNeuro at https://doi:10.18112/openneuro.ds005577.v1.0.0, reference number [ds005577]. The processed data and analysis scripts are available on OSF at https://osf.io/d8e9b/.

## Results

### Lateralization of the global ROI

#### Strength

First, we examined if there was a difference in *strength* of the LI between the two groups for each of the three tasks using the absolute LI values for the global ROIs (see Figure 2). We did not find that individuals with dyslexia had significantly lower LI values for reading, *t*(67.13) = 1.38, *P* = 0.189, rhyming, *t*(51.63) = 1.16, *P* = 0.189, or fluency, *t*(66.22) = 0.28, *P* = 0.389.

**Figure 2.**
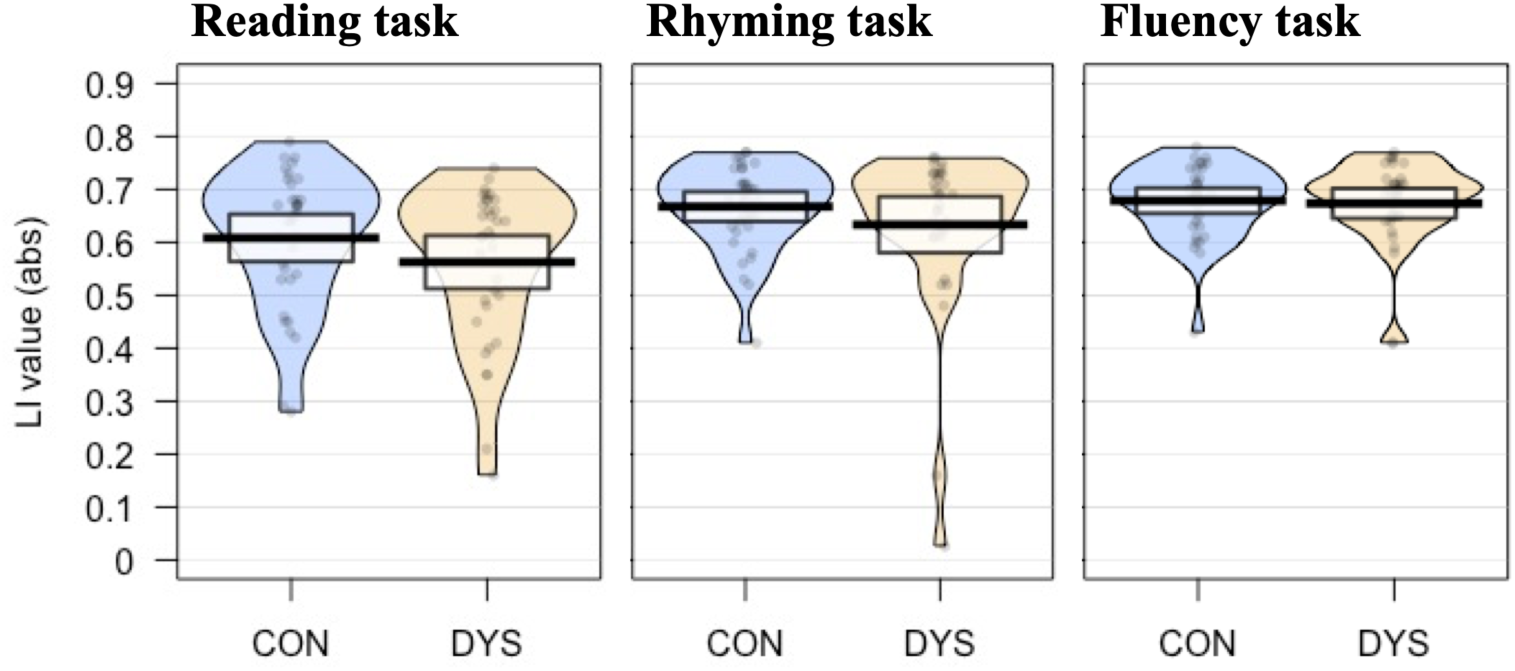
Pirate plots showing the absolute LI values for the control (n = 35) and dyslexia (n = 35) groups for the global ROI of each of the three functional MRI tasks. The bolded line shows the mean, the coloured area represents the smoothed density distribution, and the highlighted box the 95% confidence intervals. Each point represents an individual participant’s LI value. There were no significant differences in absolute LI values between the groups for any of the three tasks (reading task: *t* = 1.38, *P* = 0.189; rhyming task: *t* = 1.16, *P* = 0.189; fluency task: *t* = 0.28, *P* = 0.389). CON = control group; DYS = dyslexia group.

#### Direction

All 35 control participants showed left-hemisphere lateralization for each of the three tasks. In the dyslexia group, two participants were lateralized to the right hemisphere: one across all three tasks and one for the fluency task only, while the remaining 33 were lateralized to the left hemisphere. This difference in directional asymmetry between the two groups was not significant (*P* = 0.246 for the fluency task and *P* = 0.500 for the reading and rhyming tasks, respectively).

#### Inter-task consistency

When comparing the consistency of lateralization across the three tasks, only one participant in the dyslexia group showed inconsistent lateralization. Therefore, participants with dyslexia were no more likely than controls to have inconsistent lateralization across global ROIs for the language tasks (*P* = 0.500).

### Lateralization of the multiple task-specific ROIs

#### Strength

Here, we investigated whether there were group differences in the strength of lateralization across the series of task-specific ROIs (see Table 2). For most ROIs across all three tasks, LIs were either comparable between groups or in the unpredicted direction (i.e., higher in the dyslexic group) which meant that no statistical tests were carried out. The only exception was for LI strength in the fusiform region during the reading task, where a significant reduction was observed in the dyslexia group, *t*(58.74) = 3.19, *P* = 0.007, d = 0.76.

**Table 2.** Means, standard deviations, and statistical comparisons of global and regional LI group differences.

| Task | ROI | CON |  | DYS |  | *t*-value | df | *P*-value | effect size (*d*) | 95 % CI |
| --- | --- | --- | --- | --- | --- | --- | --- | --- | --- | --- |
|  |  | *M* | *SD* | *M* | *SD* |  |  |  |  |  |
| Read | Composite ROI | 0.61 | 0.13 | 0.56 | 0.15 | 1.38 | 67.13 | 0.189 | 0.33 | -0.01 |
|  | Fusiform gyrus | 0.54 | 0.13 | 0.41 | 0.20 | **3.19** | **58.74** | **0.007** | 0.76 | 0.06 |
|  | Inf frontal gyrus | 0.57 | 0.16 | 0.51 | 0.18 | 1.54 | 67.62 | 0.194 | 0.37 | -0.01 |
|  | Inf occipital gyrus | 0.48 | 0.22 | 0.47 | 0.16 | 0.11 | 62.09 | 0.457 | 0.03 | -0.7 |
|  | Inf parietal gyrus | 0.49 | 0.21 | 0.44 | 0.21 | 0.97 | 67.99 | 0.300 | 0.23 | -0.04 |
|  | Mid temporal gyrus | 0.55 | 0.16 | 0.52 | 0.18 | 0.85 | 67.42 | 0.300 | 0.20 | -0.03 |
|  | Precentral gyrus | 0.58 | 0.14 | 0.57 | 0.15 | 0.23 | 67.85 | 0.457 | 0.05 | -0.05 |
| Rhyme | Composite ROI | 0.67 | 0.08 | 0.63 | 0.15 | 1.16 | 51.63 | 0.189 | 0.28 | -0.02 |
|  | Cerebellum | 0.47 | 0.19 | 0.43 | 0.22 | 0.79 | 66.29 | 0.215 | 0.19 | -0.04 |
|  | Inf frontal gyrus | 0.66 | 0.08 | 0.63 | 0.12 | 1.18 | 58.34 | 0.168 | 0.28 | -0.01 |
|  | Mid temporal gyrus | 0.45 | 0.21 | 0.49 | 0.20 |  |  |  |  |  |
|  | Precentral gyrus | 0.66 | 0.06 | 0.63 | 0.13 | 1.32 | 48.49 | 0.168 | 0.32 | -0.01 |
|  | Sup motor area | 0.50 | 0.16 | 0.45 | 0.19 | 1.16 | 66.66 | 0.168 | 0.28 | -0.02 |
| Fluency | Composite ROI | 0.68 | 0.07 | 0.67 | 0.08 | 0.28 | 66.22 | 0.389 | 0.07 | -0.03 |
|  | Cerebellum | 0.57 | 0.14 | 0.56 | 0.14 | 0.25 | 68 | 0.478 | 0.06 | -0.05 |
|  | Inf frontal gyrus | 0.66 | 0.08 | 0.66 | 0.09 | 0.06 | 66.19 | 0.478 | 0.01 | -0.03 |
|  | Precentral gyrus | 0.66 | 0.12 | 0.64 | 0.09 | 0.86 | 64.49 | 0.397 | 0.21 | -0.02 |
|  | Sup motor area | 0.55 | 0.11 | 0.52 | 0.14 | 0.91 | 63.98 | 0.397 | 0.22 | -0.02 |
|  | Thalamus | 0.48 | 0.18 | 0.45 | 0.18 | 0.72 | 68.00 | 0.397 | 0.17 | -0.04 |

CON = control group; DYS = dyslexia group; M = mean; SD = standard deviation. P-values represent comparisons between the control and dyslexia groups (one-sided independent t-test). Effect sizes are reported as Cohen's *d*, and 95% confidence interval for the one-sided test. Statistical tests were not performed if the difference in means was not in the predicted direction.

#### Direction

When examining the direction of lateralization, we found that significantly more individuals with dyslexia had right-sided fusiform lateralization in the reading task compared to the control group (*P* = 0.034). No other significant differences in lateralization direction were observed (see Table 3 for further details).

**Table 3.** Distribution of left and right lateralization by ROI and group, with statistical comparisons of side of processing.

| Task | ROI | CON |  | DYS |  | Diff | P-value | 95% CI of difference |
| --- | --- | --- | --- | --- | --- | --- | --- | --- |
|  |  | Left | Right | Left | Right |  |  |  |
| Reading | Fusiform gyrus | 35 | 0 | 28 | 7 | -0.20 | **0.034** | -0.36, -0.06 |
|  | Inf frontal gyrus | 35 | 0 | 34 | 1 | -0.03 | 0.500 | -0.15, 0.07 |
|  | Inf occipital gyrus | 35 | 0 | 30 | 5 | -0.14 | 0.080 | -0.29, -0.02 |
|  | Inf parietal gyrus | 30 | 5 | 26 | 9 | -0.11 | 0.371 | -0.30, 0.08 |
|  | Mid temporal gyrus | 33 | 2 | 31 | 4 | -0.06 | 0.500 | -0.21, 0.09 |
|  | Precentral gyrus | 34 | 1 | 33 | 2 | -0.03 | 0.500 | -0.16, 0.10 |
| Rhyming | Cerebellum | 1 | 34 | 4 | 31 | -0.09 | 0.297 | -0.23, 0.05 |
|  | Inf frontal gyrus | 35 | 0 | 34 | 1 | -0.03 | 0.500 | -0.15, 0.07 |
|  | Mid temporal gyrus | 34 | 1 | 27 | 8 | -0.20 | 0.069 | -0.36, -0.04 |
|  | Precentral gyrus | 35 | 0 | 33 | 2 | -0.06 | 0.308 | -0.19, 0.05 |
|  | Sup motor area | 34 | 1 | 30 | 5 | -0.11 | 0.248 | -0.27, 0.03 |
| Fluency | Cerebellum | 35 | 0 | 34 | 1 | -0.03 | 0.500 | -0.15, 0.07 |
|  | Inf frontal gyrus | 35 | 0 | 33 | 2 | -0.06 | 0.411 | -0.19, 0.05 |
|  | Precentral | 35 | 0 | 34 | 1 | -0.03 | 0.500 | -0.15, 0.07 |
|  | Sup motor area | 35 | 0 | 33 | 2 | -0.06 | 0.411 | -0.19, 0.05 |
|  | Thalamus<sup>a</sup> | 34 | 1 | 29 | 5 | -0.12 | 0.411 | -0.27, 0.03 |

CON = control group; DYS = dyslexia group; Diff = The difference between left hemisphere processing between the two groups expressed as a proportion. *P*-values are from Fisher's Exact Tests comparing the difference in proportion between the two groups, and the 95% confidence intervals of the difference.
<sup>a</sup>One dyslexia participant lacked activation in the thalamus ROI and was excluded from that analysis

#### Inter-regional consistency

We also examined whether individuals with dyslexia had a higher incidence of inconsistent lateralization across the series of task-specific ROIs. First, we used a simple dichotomous categorization of consistent versus inconsistent lateralization. In the *reading task*, 80% of controls (28/35) and 57.14% of individuals with dyslexia (20/35) had all regions consistently lateralized to one hemisphere. This higher incidence of inconsistent lateralization among the dyslexia individuals was significant (*P* = 0.035), with 95% CIs of the difference not overlapping zero [-0.42, -0.01]. When examining only inconsistently lateralized individuals, we found that 86% (6/7) of individuals in the control group had only one deviating ROI, 14% (1/7) had two, and 0% had three deviating ROIs. Amongst dyslexia individuals, 67% (10/15) had one deviating ROI, 27% (4/15) had two, and 7% (1/15) had three deviating ROIs. A Fisher’s Exact Test indicated that the distribution of reading inconsistency ROIs did not differ significantly between the groups (*P* = 0.744).

In the *rhyming task*, 91% of controls (32/35) and 66% of dyslexia individuals (23/35) had all ROIs consistently lateralized to one hemisphere. This higher incidence of inconsistent lateralization in the dyslexia group was significant (*P* = 0.009), with the 95% CIs of the difference not overlapping zero [-0.43, -0.06]. Among the inconsistently lateralized participants, 100% (3/3) of the control group had one deviating ROI and 0% had two. In the dyslexia group, 75% (9/12) had one deviating ROI and 25% (3/12) had two. A Fisher’s Exact Test indicated that the distributions did not differ significantly between the groups (*P* = 0.484).

For the *fluency task*, 97.14% of controls (34/35) and 85.71% (30/35) of individuals with dyslexia had all ROIs consistently lateralized to one hemisphere. This difference in incidence of inconsistent lateralization was not significant (*P* = 0.099) with the 95% CIs of the difference overlapping zero [-0.27, 0.03]. Among the inconsistently lateralized individuals, 100% (1/1) of the control participants had only one deviating ROI. Among dyslexia individuals, 80% (4/5) had one deviating ROI and 20% (1/5) had two. This difference in frequencies was also not significant (*P* = 0.833).

### Reading performance across consistency groups

Here, we examined reading performance in individuals with dyslexia with consistent versus inconsistent lateralization across regions in the *reading task*. When comparing the consistently lateralized group (*n* = 15, *M* = 75.55, *SD* = 13.49) with the inconsistently lateralized group (*n* = 20, *M* = 67.60, *SD* = 18.48), we did not find that reading performance was significantly worse in the inconsistently lateralized group, *t*(24.54) = 1.41, *P* = 0.086, *d* = 0.50, and the 95% CI of the difference overlapped with zero [-1.70] (see Figure 3).

**Figure 3.**
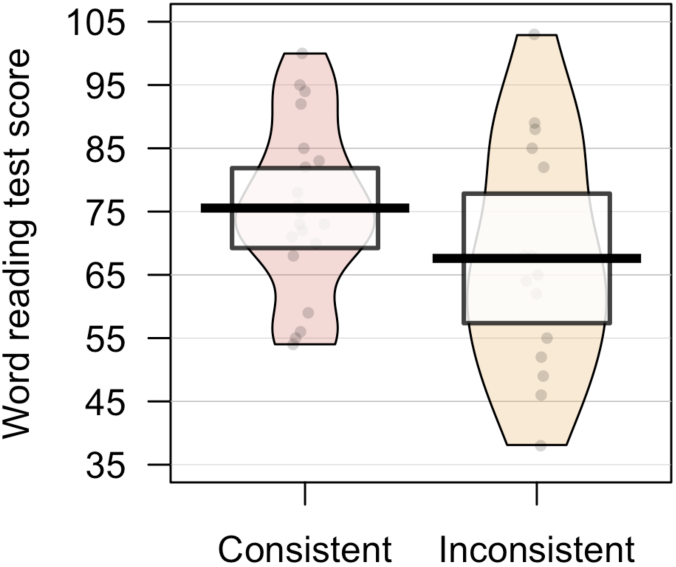
Word reading test scores for dyslexia participants with consistent (n = 20) versus inconsistent (n = 15) reading activation patterns. The bolded line shows the mean, the coloured area represents the smoothed density distribution, and the highlighted box the 95% confidence intervals. Each point represents an individual participant’s word reading test score. The inconsistently lateralized group did not have a significantly lower word reading test score compared to consistently lateralized individuals (*t* = 1.41, *P* = 0.086).

### Follow-up analyses

#### Fusiform lateralization within the dyslexia group

We further wanted to explore the functional relevance of the fusiform gyrus for reading by examining its lateralization within the *reading task*. When comparing the absolute strength of lateralization between the consistently lateralized group (*n* = 20, *M* = 0.49, *SD* = 0.18) and the inconsistently lateralized group (*n* = 15, *M* = 0.31, *SD* = 0.18), we found that lateralization strength in the fusiform gyrus was significantly lower in the inconsistently lateralized group, *t*(30.23) = 2.84*, P* = .004, d = 0.97, 95% CI of the difference did not overlap zero [0.07] (see Figure 4). There was no significant relationship between the absolute LI strength of the fusiform gyrus and reading test score, *r*(33)= 0.13, *P* = 0.440.

**Figure 4.**
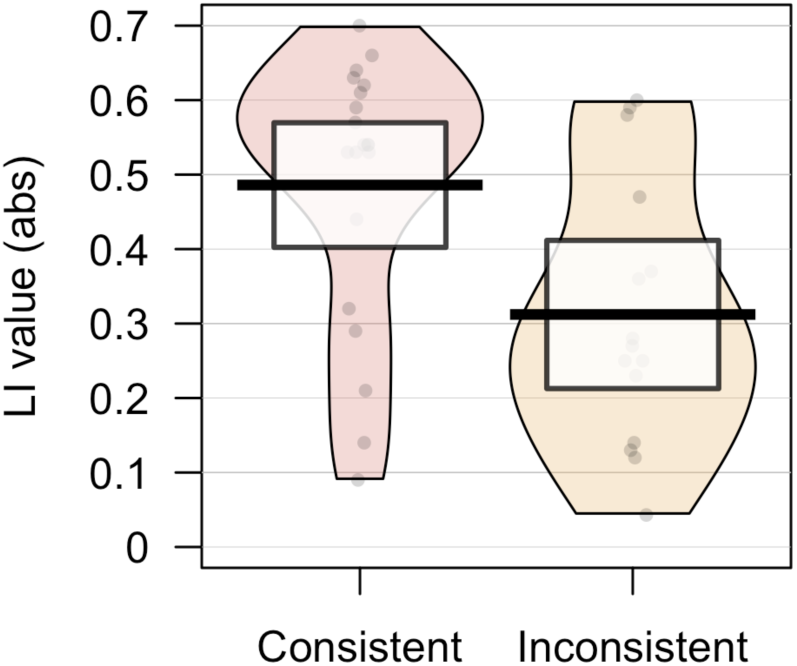
Absolute LI values in the fusiform ROI for consistently (n = 20) and inconsistently (n = 15) lateralized individuals in the dyslexia group. The bolded line shows the mean, the coloured area represents the smoothed density distribution, and the highlighted box the 95% confidence intervals. Each point represents an individual participant’s absolute LI value. The bolded line shows the mean and the highlighted area the 95% confidence intervals. The group of inconsistently lateralized individuals had significantly lower absolute LI values compared to the consistently lateralized group (*t* = 2.84*, P* = .004).

## Discussion

Despite a century of research, it remains unclear whether and how hemispheric asymmetries for language are altered in dyslexia. Addressing limitations of previous studies, we conducted the most in-depth and comprehensive investigation of functional hemispheric asymmetry in dyslexia to date, accounting for individual differences. Across three distinct language tasks, we tease apart degree and direction of asymmetry while also examining between-task and between-region consistency in language lateralization. Our findings are not only important for understanding dyslexia but also for informing the broader investigation of hemispheric asymmetry, with several noteworthy outcomes.

First, these data do not support our hypothesis that individuals with dyslexia have generally weaker lateralization compared to a control group without reading difficulties. Relative to controls, participants with dyslexia demonstrated comparable lateralization across almost all ROIs in every language task, both at the global and regional levels. The only exception was for the fusiform region in the reading task, in which individuals with dyslexia were on average less lateralized. This contrasts with previous studies reporting decreased lateralization in dyslexic groups in areas beyond the fusiform gyrus and using different task types (e.g., ^31,37^). Unlike earlier studies, we examined asymmetry using absolute values, enabling us to separate direction (left vs. right) from degree of lateralization. It is likely that some results in previous studies have been skewed by instances of right-sided values, which can substantially affect lateralization metrics, especially in small samples (to illustrate this further, see Supplementary Table 2, where we calculate asymmetries for each ROI/task using original values).

Although we observed a higher incidence of right hemisphere lateralization among individuals with dyslexia, this difference reached significance only in the fusiform gyrus during the reading task. This region encompasses the so-called visual word form area (VWFA), which is crucial for fluent word reading. Hypoactivity in the left VWFA is one of the most consistent findings in group-averaged activation maps of dyslexia.^24,25,65^ Although our between-group comparisons did not reveal significant differences in VWFA activation, the dyslexia group showed, on average, decreased lateralization in this area. This group difference also remained in post-hoc exploratory analyses when only individuals with left-sided fusiform activation were included (CON n = 35, DYS n = 28; *t*(45.39) = 2.21, *P* = 0.016, one-sided). Thus, our analyses indicate reduced hemispheric asymmetry in the dyslexia group, but not necessarily consistent hypoactivation within this region.

Bradshaw et al^39^ found that adults with DLD showed significantly higher rates of inconsistent lateralization across different language tasks compared to a control group. Here, we did not find that inconsistent lateralization *across tasks* was more common among individuals with dyslexia. Although only two out of three tasks were the same for our and their study, the diverging results are more likely due to methodological differences. The group of Bradshaw et al. was a heterogenous group of diagnoses of developmental disorders affecting language and communication, including dyslexia, autism and dyspraxia. It could be that different language disorders are associated with different lateralization profiles depending on the specific impairment. In addition, the use of fTCD meant that their lateralization measures were restricted to covering the middle cerebral artery territory globally, whilst we were able to examine a specific selection of language-related regions. It is unclear how compensatory mechanisms affect the LI obtained from fTCD and the authors speculated that their right-sided LI values could have been the result of increased recruitment of additional right hemisphere resources to compensate for deficient left hemisphere processing.

In our data, we instead observed a higher incidence of inconsistent lateralization across task-specific regions in individuals with dyslexia. Interestingly, this inconsistency was apparent only in reading and phonological processing when comparing the number of consistent versus inconsistent individuals between groups. Verbal fluency, however, showed comparable patterns in both groups. Of course, not every single person in the dyslexia group showed inconsistent lateralization. It could be that other neural regions, beyond those chosen for our analysis, may be important. Or it could be that not all individuals with dyslexia have an inconsistently lateralized language system, highlighting the heterogeneity of developmental disorders. Alternatively, these findings suggest that inconsistent lateralization may be related to specific symptoms of dyslexia. It has been proposed that dyslexia consists of multiple subtypes with differing cognitive and reading-related profiles.^66–69^ This heterogeneity in symptom profiles, which may rely on different underlying mechanisms, could contribute to the variation in asymmetry patterns observed in our dyslexia sample. Overall, our findings support the view that dyslexia may stem from diverse neurobiological causes, with atypical lateralization possibly serving as one of several risk factors.

More importantly, the differences in lateralization consistency between dyslexic and control participants suggest that stable hemispheric dominance, rather than direction, may be crucial for efficient language processing. Kosslyn (1987), for example, proposed that an intrinsic leftward bias in speech control creates a cascading effect, leading other language functions to lateralize similarly, thus fostering efficiency. When individuals lack this consistency bias, they may have more diffused and less efficient neural networks across both hemispheres, potentially resulting in impaired language abilities.^70^ Similarly, Bishop et al. suggested that inconsistency in lateralization (albeit across language functions) may result in a less efficient language network.^38^ In the current study, inconsistent lateralization was only observed across different regions within tasks, and only for reading and rhyming. These findings underscore the potential role of lateralization patterns as foundational to cognitive performance in reading and language skills. Of course, we can only say that there is an association between inconsistent lateralisation and dyslexia. From our data, we cannot explain the origin of that relationship. It could be that inconsistent lateralization is a cause of the symptoms seen in dyslexia. Alternatively, it could be that it is the consequence of dyslexia, if dyslexia has an influence on how the brain develops or they could co-occur because they have a common origin, but are not directly caused by one another (for an excellent discussion see ^71^).

Our findings also have broader implications for how hemispheric asymmetry is conceptualized in research. Traditionally, asymmetry is treated simply as a left-right dominance issue or, alternatively, as the absence of asymmetry. However, our study demonstrates that dyslexia is not characterized by a lack of asymmetrical processing. When we calculate a global laterality index, both groups show similar consistency and strength of lateralization, suggesting no apparent difference. At first glance, these traditional metrics might lead to the conclusion that dyslexic and control groups have comparable lateralization patterns. However, by examining lateralization in a more nuanced way - looking at task-specific regions and consistency within tasks - we uncover significant differences. These findings suggest that hemispheric asymmetry is not just about whether the left or right hemisphere is dominant. Instead, it also encompasses the stability and precision of how lateralized processing occurs across and within various neural networks.

One limitation of our study is that the classification of hemispheric dominance depends on the method used to calculate the laterality index (LI). We used the LI toolbox,^52^ which is widely regarded as the current gold standard^27^. A criticism of this method is that it can produce wide confidence intervals, since it estimates LIs across multiple thresholds guided by the maximum *t*-value in the data. While estimates are typically consistent within thresholds, they may vary across them, resulting in broader intervals.^28^ At the same time, this approach reduces reliance on any single, potentially arbitrary cut-off and has been shown to produce reliable LI values across repeated sessions.^72^ Thus, while the method may yield wider confidence intervals, we consider the resulting estimates more robust and less susceptible to arbitrary thresholding. An alternative approach is the mirror method proposed by Bishop et al.^28^ which instead estimates the LI using a threshold-free bootstrapping procedure.

## Conclusions

In conclusion, our findings challenge longstanding assumptions about hemispheric asymmetry in dyslexia by demonstrating that this condition is not characterized by a general weaker or absent lateralization, but rather by inconsistent and regionally atypical patterns. While global lateralization indices reveal similarities between dyslexic and control groups, a closer examination highlights significant differences in the stability and specificity of lateralized processing across language-related regions. These inconsistencies, particularly in reading and phonological tasks, suggest that stable hemispheric dominance may be more critical for efficient language processing than the direction of lateralization. This nuanced perspective not only advances our understanding of the neurobiological underpinnings of dyslexia but also underscores the importance of examining lateralization beyond traditional metrics. Future research should explore whether these inconsistencies are a cause or consequence of dyslexia or if they arise from shared developmental origins. By adopting more detailed and region-specific analyses, researchers can further elucidate the role of hemispheric asymmetry in language disorders and contribute to more targeted interventions for individuals with dyslexia.

## Acknowledgements

We would like to thank the participants who took part in this study.

## Supplementary material

Supplementary material is available at *Brain* online.

