## Supplemental materials for "Revisiting atypical language lateralization in dyslexia"

Supplement to manuscript “*Revisiting atypical language lateralization in dyslexia*”.

**A. Group-level and between-group fMRI activations during the reading task**

|  |  | MNI coordinates (peak voxel) | | |  |  |
| --- | --- | --- | --- | --- | --- | --- |
| Contrast | Anatomical definition | *X* | *Y* | *Z* | *t-value* | *Cluster size* |
| Control group | **Left precentral gyrus** | **-48** | **10** | **30** | **9.46** | **2966** |
|  | Left inferior frontal gyrus, opercular part | -48 | 16 | 10 | 9.34 |  |
|  | Left inferior frontal gyrus, triangular part | -28 | 30 | 0 | 8.81 |  |
|  | **Left inferior occipital gyrus** | **-26** | **-100** | **-10** | **9.46** | **1457** |
|  | Left fusiform gyrus | -36 | -47 | -18 | 8.7 |  |
|  | Left inferior temporal gyrus | -48 | -57 | -13 | 8.37 |  |
|  | **Right lobule VIIB of cerebellar hemisphere** | **19** | **-77** | **-48** | **9.07** | **1020** |
|  | Right lobule VIII of cerebellar hemisphere | 26 | -72 | -50 | 7.77 |  |
|  | Right lobule VI of cerebellar hemisphere | 9 | -77 | -20 | 6.8 |  |
|  | **Left superior frontal gyrus, dorsolateral** | **-11** | **13** | **54** | **8.63** | **1088** |
|  | Right superior frontal gyrus, medial | 9 | 20 | 42 | 8.2 |  |
|  | Left supplementary motor area | -4 | 10 | 57 | 8.06 |  |
|  | **Right insula** | **32** | **23** | **-3** | **8.28** | **1019** |
|  | Right inferior frontal gyrus, pars orbitalis | 42 | 26 | -10 | 7.33 |  |
|  | Right inferior frontal gyrus, triangular part | 46 | 16 | 22 | 5.87 |  |
|  | **Right inferior frontal gyrus, triangular part** | **24** | **-94** | **-6** | **7.89** | **153** |
|  | Right inferior occipital gyrus | 42 | -82 | -10 | 3.89 |  |
|  | **Left middle temporal gyrus** | **-54** | **-40** | **7** | **7.71** | **240** |
|  | Left middle temporal gyrus | -44 | -42 | 4 | 4.63 |  |
|  | Left middle temporal gyrus | -56 | -60 | 7 | 4.28 |  |
|  | **Left anterior cingulate cortex, supracallosal** | **-6** | **3** | **30** | **6.92** | **58** |
|  | **Left middle occipital gyrus** | **-28** | **-70** | **40** | **6.66** | **77** |
|  | **Right calcarine fissure and surrounding cortex** | **16** | **-64** | **12** | **5.63** | **510** |
|  | Left calcarine fissure and surrounding cortex | -16 | -77 | 14 | 5.49 |  |
|  | Right calcarine fissure and surrounding cortex | 19 | -67 | 4 | 5.05 |  |
|  | **Left caudate nucleus** | **-18** | **-12** | **20** | **5.37** | **86** |
|  | Left mediodorsal lateral parvocellular | -8 | -12 | 7 | 4.96 |  |
| Dyslexia group | **Left inferior frontal gyrus, triangular part** | **-34** | **26** | **-3** | **10.33** | **3292** |
|  | Left inferior frontal gyrus, opercular part | -41 | 8 | 27 | 9.3 |  |
|  | Left temporal pole | -48 | 13 | -13 | 8 |  |
|  | **Right insula** | **39** | **26** | **-3** | **9.77** | **742** |
|  | Right insula | 32 | 26 | 2 | 9.62 |  |
|  | Right inferior frontal gyrus, pars orbitalis | 36 | 36 | -6 | 6.09 |  |
|  | **Left inferior occipital gyrus** | **-24** | **-97** | **-8** | **9.65** | **2249** |
|  | Right calcarine fissure and surrounding cortex | 19 | -97 | -6 | 9.61 |  |
|  | Left inferior occipital gyrus | -41 | -77 | -16 | 8.55 |  |
|  | **Left supplementary motor area** | **-1** | **18** | **50** | **9.38** | **927** |
|  | Right middle cingulate & paracingulate gyri | 12 | 28 | 32 | 6.93 |  |
|  | Left supplementary motor area | -1 | 0 | 62 | 6.58 |  |
|  | **Right lobule VI of cerebellar hemisphere** | **29** | **-67** | **-23** | **8.24** | **1338** |
|  | Right lobule VIII of cerebellar hemisphere | 29 | -67 | -56 | 8.07 |  |
|  | Right lobule VIII of cerebellar hemisphere | 34 | -62 | -50 | 8.01 |  |
|  | **Left middle temporal gyrus** | **-51** | **-40** | **7** | **5.59** | **123** |
|  | Left middle temporal gyrus | -64 | -32 | 4 |  |  |
| CON>DYS | **Left supramarginal gyrus** | **-66** | **-27** | **37** | **4.43** | **88** |
|  | Left supramarginal gyrus | -64 | -34 | 44 | 4.23 |  |
|  | Left supramarginal gyrus | -66 | -30 | 27 | 4.08 |  |
| DYS>CON | **Lobule VI of right cerebellar hemisphere** | **26** | **-62** | **-23** | **4.77** | **64** |

**Supplementary Table 1.** Whole-brain analysis results for the contrast reading > control in the control group (CON) and the dyslexia group (DYS), as well as between-group comparisons (CON > DYS and DYS > CON). Activation patterns were identified at a voxel-level threshold of p < .001 (uncorrected) with cluster-level FWE correction at p < .05.

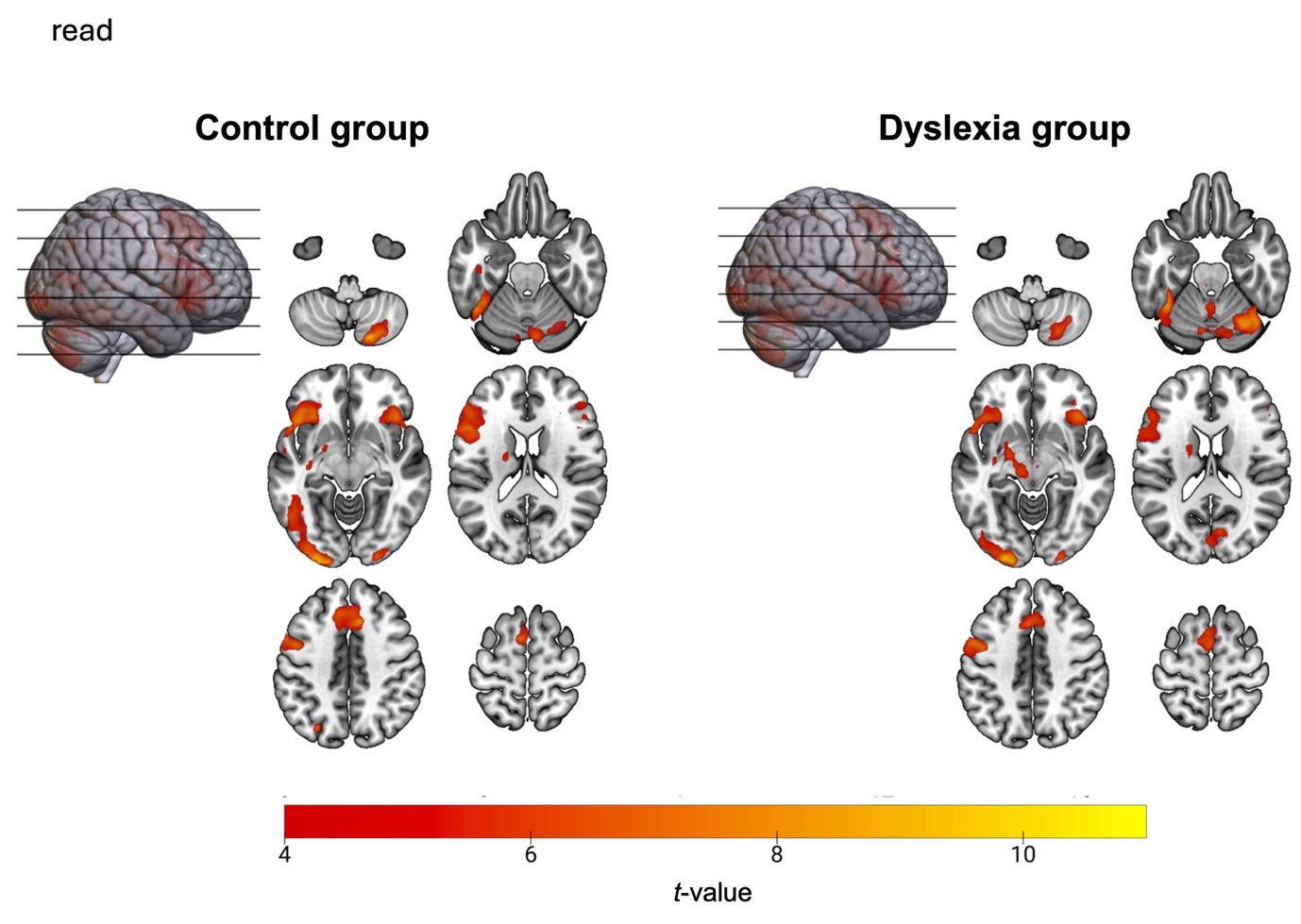

**Supplementary Figure 1.** Significant fMRI activations during the reading task shown separately for the control group and the dyslexia group, voxel-level threshold: p < 0.001 uncorrected; cluster-level threshold: p < 0.05 FWE-corrected.

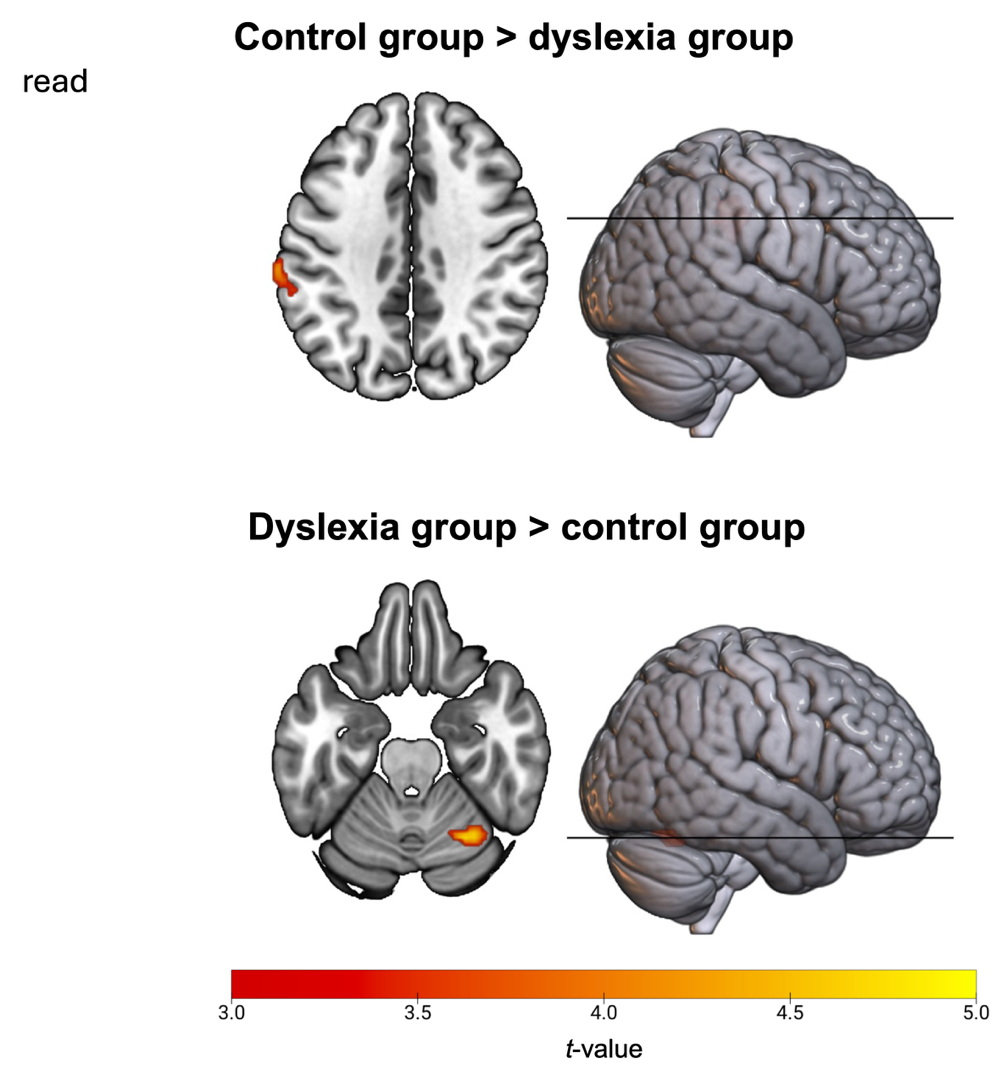

**Supplementary Figure 2.** Significant between-group fMRI activations during the reading task (controls > dyslexia and dyslexia > controls), voxel-level threshold: p < 0.001 uncorrected, cluster-level threshold: p < 0.05 FWE-corrected.

**B. Group-level and between-group fMRI activations during the rhyming task**

|  |  | MNI coordinates (peak voxel) | | |  |  |
| --- | --- | --- | --- | --- | --- | --- |
| **Contrast** | **Anatomical definition** | ***X*** | ***Y*** | ***Z*** | ***t-value*** | ***Cluster size*** |
| **Controls** | **Left inferior temporal gyrus** | **-48** | **-64** | **-8** | **15.78** | **24308** |
|  | Right lobule VIII of cerebellar hemisphere | 32 | -70 | -48 | 15.15 |  |
|  | Right lobule VI of cerebellar hemisphere | 9 | -74 | -20 | 15.03 |  |
|  | **Left supplementary motor area** | **-6** | **8** | **60** | **14.58** | **1683** |
|  | Right middle cingulate & paracingulate gyri | 12 | 33 | 30 | 11.11 |  |
|  | Left superior frontal gyrus, dorsolateral | -11 | 13 | 50 | 10.96 |  |
|  | **Left superior temporal gyrus** | **-51** | **-44** | **12** | **7.77** | **242** |
|  | Left middle temporal gyrus | -56 | -30 | 2 | 5.53 |  |
|  | Left supramarginal gyrus | -56 | -44 | 24 | 4.19 |  |
|  | **Left calcarine fissure and surrounding cortex** | **-16** | **-80** | **10** | **4.54** | **87** |
|  | Left calcarine fissure and surrounding cortex | -11 | -70 | 10 | 4.28 |  |
| **DYS** | **Left inferior frontal gyrus, triangular part** | **-28** | **30** | **2** | **14.65** | **16483** |
|  | Right Insula | 34 | 26 | -6 | 13.64 |  |
|  | Left red nucleus | -6 | -27 | -13 | 13.17 |  |
|  | **Left supplementary motor area** | **-1** | **20** | **47** | **11.29** | **993** |
|  | Left supplementary motor area | -4 | 10 | 52 | 10.42 |  |
|  | Right superior frontal gyrus, medial | 6 | 20 | 42 | 10.36 |  |
|  | **Right lobule X of cerebellar hemisphere** | **22** | **-40** | **-43** | **9.39** | **64** |
|  | **Left lobule VIII of cerebellar hemisphere** | **-31** | **-67** | **-53** | **5.58** | **83** |
|  | Left lobule VIIB of cerebellar hemisphere | -38 | -62 | -53 | 5.00 |  |
|  | **Left middle temporal gyrus** | **-56** | **-30** | **2** | **5.39** | **58** |
|  | Left middle temporal gyrus | -54 | -40 | 4 | 4.26 |  |
| **CON>DYS** | **Left precentral gyrus** | **-36** | **8** | **37** | **5.28** | **74** |
|  | Left middle frontal gyrus | -34 | 16 | 40 | 3.83 |  |
|  | **Left superior frontal gyrus, medial** | **-8** | **30** | **50** | **4.63** | **86** |
|  | Left superior frontal gyrus, dorsolateral | -14 | 28 | 34 | 3.77 |  |
|  | **Left middle occipital gyrus** | **-28** | **-74** | **22** | **4.18** | **59** |
|  | Left middle occipital gyrus | -26 | -72 | 30 | 3.85 |  |
| **DYS>CON** |  | **-** | **-** | **-** | **-** | **-** |

**Supplementary Table 1.** Whole-brain analysis results for the contrast rhyming > control in the control group (CON) and the dyslexia group (DYS), as well as between-group comparisons (CON > DYS and DYS > CON). Activation patterns were identified at a voxel-level threshold of p < .001 (uncorrected) with cluster-level FWE correction at p < .05.

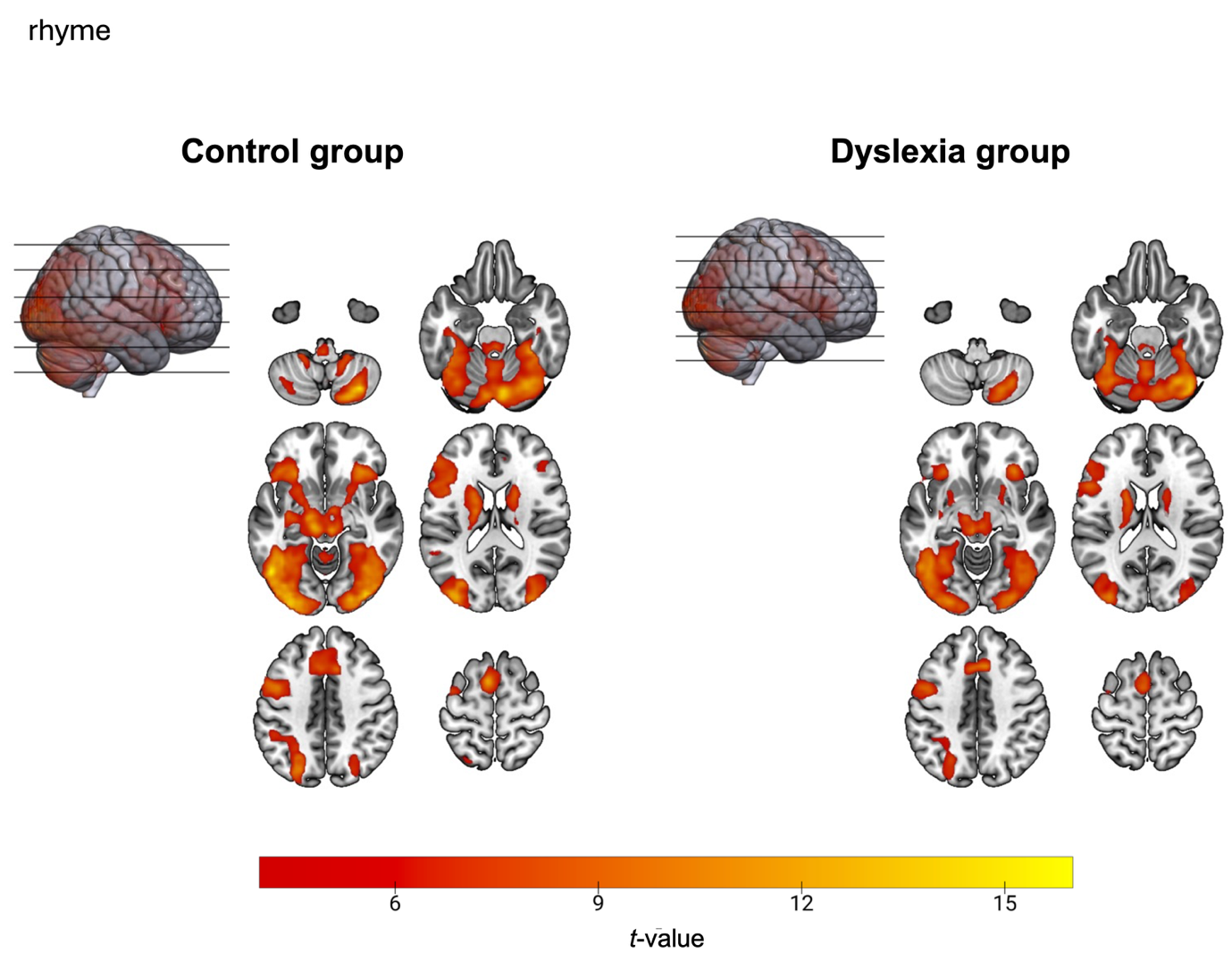

**Supplementary Figure 3.** Significant fMRI activations during the rhyming task shown separately for the control group and the dyslexia group, voxel-level threshold: p < 0.001 uncorrected; cluster-level threshold: p < 0.05 FWE-corrected.

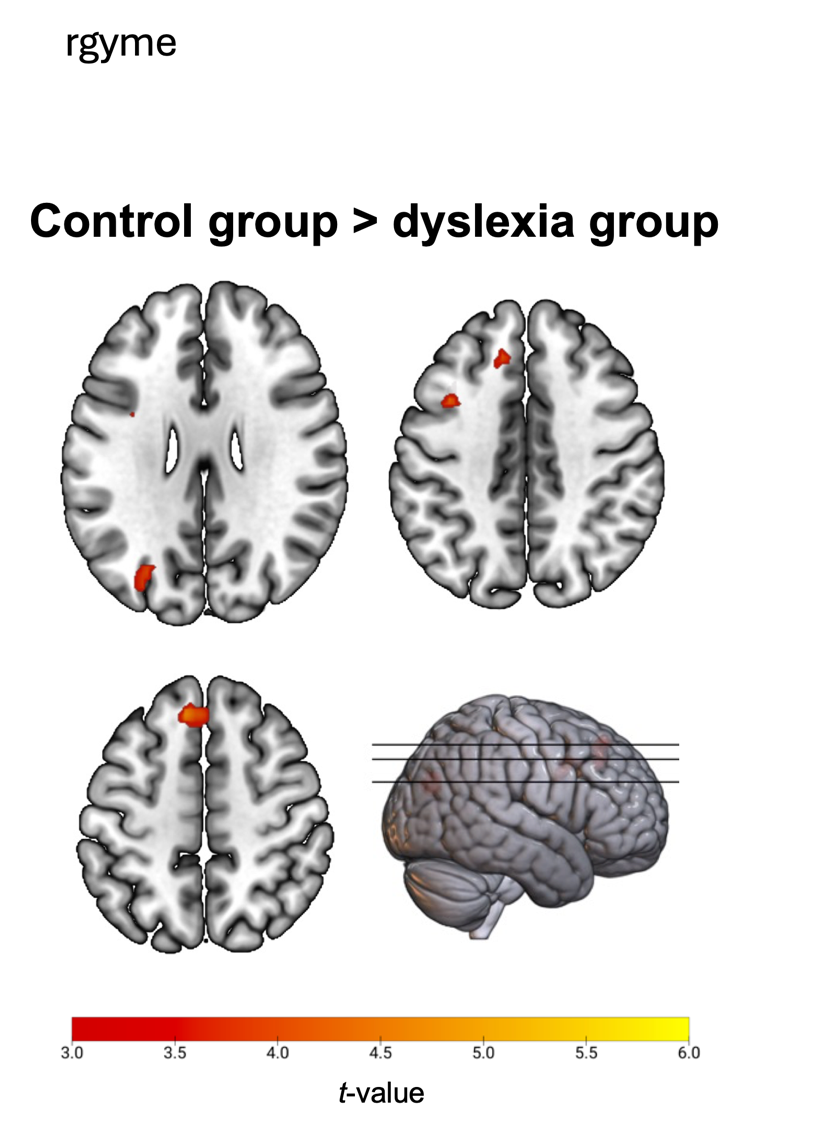

**Supplementary Figure 4.** Significant between-group fMRI activations during the rhyming task for controls > dyslexia with voxel-level threshold: p < 0.001 uncorrected, cluster-level threshold: p < 0.05 FWE-corrected.

**C. Group-level and between-group fMRI activations during the fluency task**

|  |  | MNI coordinates (peak voxel) | | |  |  |
| --- | --- | --- | --- | --- | --- | --- |
| **Contrast** | **Anatomical definition** | ***X*** | ***Y*** | ***Z*** | ***t-value*** | ***Cluster size*** |
| **Wordgen Control** | **Right lobule VI of cerebellar hemisphere** | **26** | **-62** | **-26** | **16.06** | **18154** |
|  | Right crus I of cerebellar hemisphere | 44 | -60 | -30 | 15.66 |  |
|  | Left supplementary motor area | -6 | 8 | 60 | 15.5 |  |
|  | **Left inferior parietal gyrus** | **-48** | **-40** | **47** | **9.59** | **1082** |
|  | Left superior parietal gyrus | -28 | -62 | 47 | 8.64 |  |
|  | Left middle occipital gyrus | -24 | -72 | 30 | 7.71 |  |
|  | **Left lobule IX of cerebellar hemisphere** | **-6** | **-37** | **-43** | **7.59** | **184** |
|  | Left lobule X of cerebellar hemisphere | -1 | -30 | -48 | 6.82 |  |
|  | Left lobule IX of cerebellar hemisphere | -4 | -44 | -56 | 5.57 |  |
|  | **Right middle frontal gyrus** | **39** | **46** | **20** | **5.66** | **147** |
|  | Right middle frontal gyrus | 32 | 43 | 12 | 5.28 |  |
|  | Right middle frontal gyrus | 32 | 30 | 22 | 4.34 |  |
|  | **Right angular gyrus** | **29** | **-50** | **32** | **5.48** | **69** |
|  | Right precuneus | 22 | -50 | 44 | 4.65 |  |
|  | Right angular gyrus | 32 | -60 | 40 | 3.91 |  |
| **Wordgen Dyslexia** |  |  |  |  |  |  |
|  | **Right lobule VIII of cerebellar hemisphere** | **29** | **-67** | **-56** | **15.09** | **1377** |
|  | Right crus I of cerebellar hemisphere | 44 | -62 | -28 | 13.59 |  |
|  | Right lobule VI of cerebellar hemisphere | 26 | -64 | -23 | 12.87 |  |
|  | **Left insula** | **-31** | **26** | **2** | **14.67** | **5913** |
|  | Right insula | 32 | 23 | 2 | 12.71 |  |
|  | Left inferior frontal gyrus, pars orbitalis | -41 | 26 | -6 | 11.04 |  |
|  | **Left supplementary motor area** | **-4** | **16** | **44** | **12.62** | **2007** |
|  | Left superior frontal gyrus, medial | -4 | 23 | 37 | 12.11 |  |
|  | Left supplementary motor area | -1 | 8 | 60 | 12.09 |  |
|  | **Left crus I of cerebellar hemisphere** | **-48** | **-60** | **-30** | **6.55** | **189** |
|  | Left lobule VI of cerebellar hemisphere | -34 | -54 | -30 | 6.47 |  |
|  | **Left middle occipital gyrus** | **-28** | **-67** | **40** | **6.4** | **222** |
|  | Left inferior parietal gyrus | -28 | -60 | 44 | 5.21 |  |
|  | Left angular gyrus | -28 | -54 | 30 | 4.86 |  |
|  | **Left middle temporal gyrus** | **-61** | **-32** | **4** | **6.37** | **104** |
|  | **Left inferior parietal gyrus** | **-44** | **-40** | **42** | **5.85** | **96** |
|  | Left inferior parietal gyrus | -51 | -34 | 44 | 4.77 |  |
|  | **Left putamen** | **-34** | **-20** | **-8** | **5.72** | **106** |
|  | Left hippocampus | -28 | -34 | 7 | 5.38 |  |
|  | Left hippocampus | -34 | -32 | 0 | 4.44 |  |
|  | **Right calcarine fissure and surrounding cortex** | **32** | **-52** | **7** | **5.45** | **68** |
|  | Right hippocampus | 34 | -44 | 4 | 4.3 |  |
|  | Right hippocampus | 34 | -34 | 0 | 4.02 |  |
|  | **Left middle frontal gyrus** | **-31** | **46** | **20** | **5.15** | **81** |
|  | Left middle frontal gyrus | -36 | 53 | 20 | 4.49 |  |
| **CON>DYS** | **Right lobule III of cerebellar hemisphere** | **14** | **-37** | **-23** | **5.44** | **93** |
|  | Right lobule IX of cerebellar hemisphere | 14 | -50 | -30 | 4.33 |  |
|  | Right lobule IV, V of cerebellar hemisphere | 22 | -37 | -28 | 4.25 |  |
|  | **Right precentral gyrus** | **22** | **-4** | **47** | **4.74** | **65** |
|  | Right precentral gyrus | 32 | -10 | 54 | 3.76 |  |
|  | **Left middle frontal gyrus** | **-31** | **3** | **34** | **4.62** | **129** |
|  | Left precentral gyrus | -46 | 0 | 20 | 4.26 |  |
|  | Left precentral gyrus | -46 | 6 | 34 | 4.22 |  |
|  | **Right angular gyrus** | **24** | **-50** | **42** | **4.38** | **93** |
|  | Right angular gyrus | 29 | -50 | 34 | 4.21 |  |
|  | Right inferior parietal gyrus | 36 | -42 | 47 | 4.12 |  |
| **DYS>CON** |  | **-** | **-** | **-** | **-** | **-** |

**Supplementary Table 3.** Whole-brain analysis results for the contrast fluency > control in the control group (CON) and the dyslexia group (DYS), as well as between-group comparisons (CON > DYS and DYS > CON). Activation patterns were identified at a voxel-level threshold of p < .001 (uncorrected) with cluster-level FWE correction at p < .05.

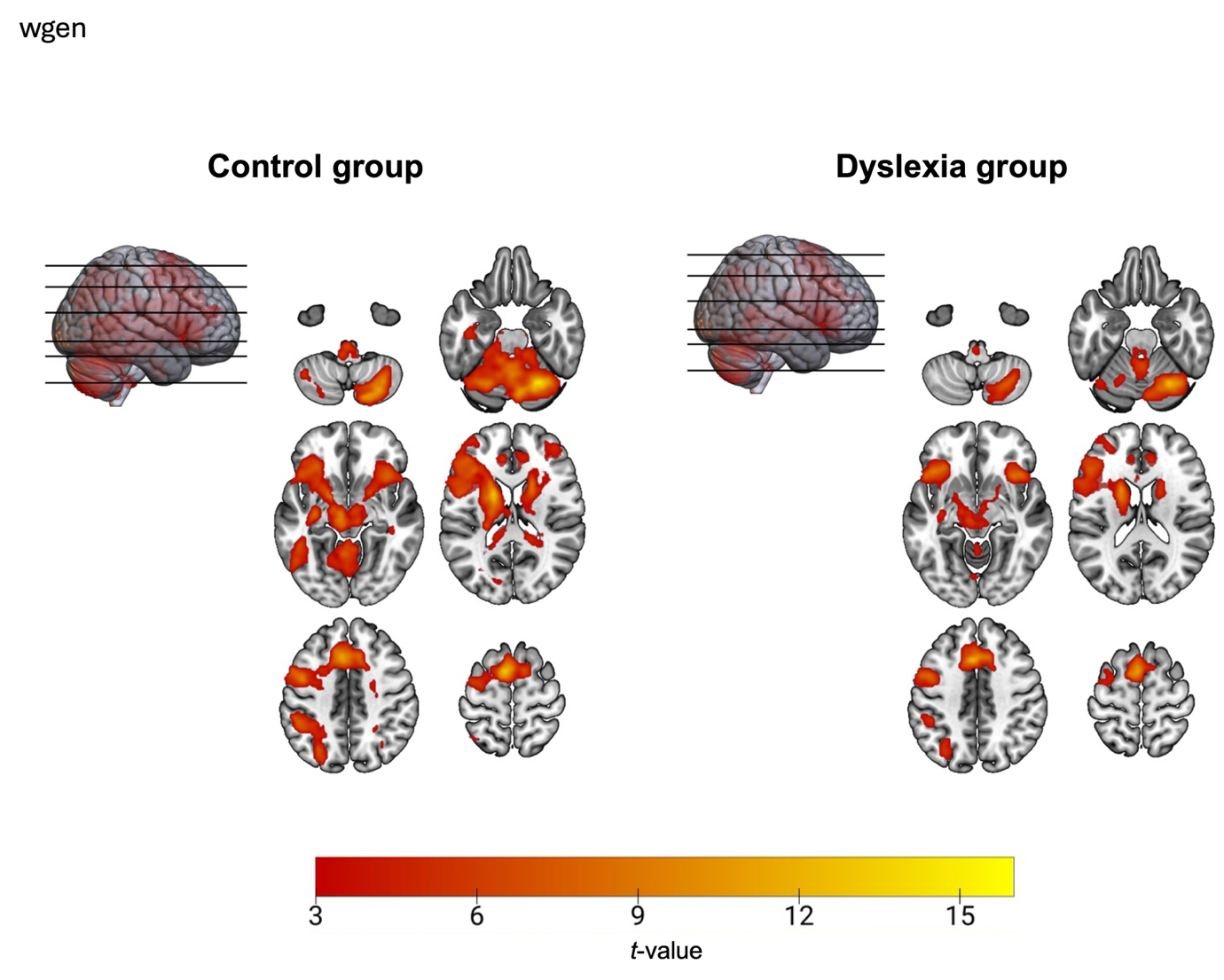

**Supplementary Figure 5.** Significant fMRI activations during the fluency task shown separately for the control group and the dyslexia group, voxel-level threshold: p < 0.001 uncorrected; cluster-level threshold: p < 0.05 FWE-corrected.

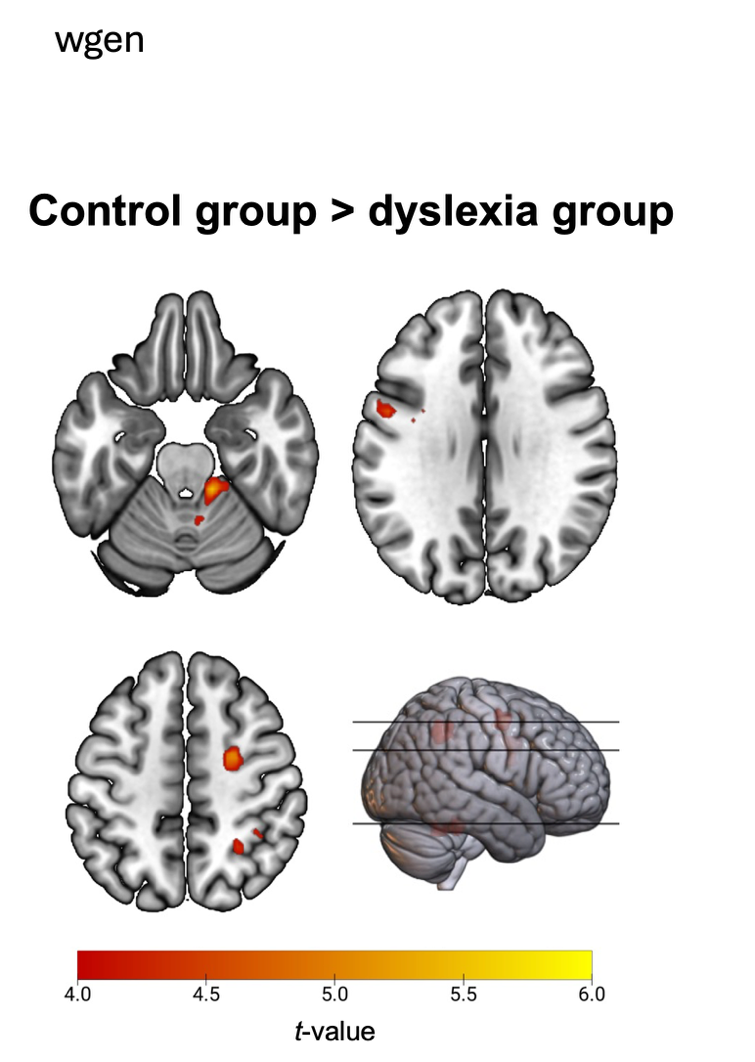

**Supplementary Figure 6**. Significant between-group fMRI activations during the fluency task for controls > dyslexia with voxel-level threshold: p < 0.001 uncorrected, cluster-level threshold: p < 0.05 FWE-corrected.

**D. Mean laterality indices (LIs), standard deviations, and one-sample tests against zero for each ROI**

| **Task** | **ROI** | **Group** | **Mean LI** | **SD** | ***t*-value** | ***P*-value** |
| --- | --- | --- | --- | --- | --- | --- |
| **Reading task** | Composite ROI | CON | -0.61 | 0.13 | -27.67 | **< 0.001** |
|  |  | DYS | -0.52 | 0.25 | -12.18 | **< 0.001** |
|  | Fusiform gyrus | CON | -0.54 | 0.13 | -24.64 | **< 0.001** |
|  |  | DYS | -0.30 | 0.35 | -5.13 | **< 0.001** |
|  | Inf frontal gyrus | CON | -0.57 | 0.16 | -20.75 | **< 0.001** |
|  |  | DYS | -0.47 | 0.26 | -10.50 | **< 0.001** |
|  | Inf occipital gyrus | CON | -0.48 | 0.22 | -12.64 | **< 0.001** |
|  |  | DYS | -0.34 | 0.37 | -5.45 | **< 0.001** |
|  | Inf parietal gyrus | CON | -0.42 | 0.33 | -7.56 | **< 0.001** |
|  |  | DYS | -0.24 | 0.43 | -3.33 | **0.002** |
|  | Mid temporal gyrus | CON | -0.52 | 0.25 | -12.29 | **< 0.001** |
|  |  | DYS | -0.45 | 0.32 | -8.18 | **< 0.001** |
|  | Precentral gyrus | CON | -0.57 | 0.16 | -20.79 | **< 0.001** |
|  |  | DYS | -0.55 | 0.22 | -14.64 | **< 0.001** |
| **Rhyming task** | Composite ROI | CON | -0.67 | 0.08 | -48.45 | **< 0.001** |
|  |  | DYS | -0.59 | 0.28 | -12.67 | **< 0.001** |
|  | Cerebellum | CON | 0.44 | 0.25 | 10.54 | **< 0.001** |
|  |  | DYS | 0.38 | 0.30 | 7.43 | **< 0.001** |
|  | Inf frontal gyrus | CON | -0.66 | 0.08 | -49.60 | **< 0.001** |
|  |  | DYS | -0.59 | 0.26 | -13.65 | **< 0.001** |
|  | Mid temporal gyrus | CON | -0.45 | 0.22 | -11.84 | **< 0.001** |
|  |  | DYS | -0.36 | 0.39 | -5.50 | **< 0.001** |
|  | Precentral gyrus | CON | -0.66 | 0.06 | -62.56 | **< 0.001** |
|  |  | DYS | -0.58 | 0.27 | -12.76 | **< 0.001** |
|  | Sup motor area | CON | -0.49 | 0.20 | -14.67 | **< 0.001** |
|  |  | DYS | -0.37 | 0.32 | -6.91 | **< 0.001** |
| **Fluency task** | Composite ROI | CON | -0.68 | 0.07 | -57.59 | **< 0.001** |
|  |  | DYS | -0.61 | 0.30 | -12.05 | **< 0.001** |
|  | Cerebellum | CON | 0.57 | 0.14 | 23.38 | **< 0.001** |
|  |  | DYS | 0.52 | 0.25 | 12.19 | **< 0.001** |
|  | Inf frontal gyrus | CON | -0.66 | 0.08 | -49.81 | **< 0.001** |
|  |  | DYS | -0.60 | 0.31 | -11.41 | **< 0.001** |
|  | Precentral gyrus | CON | -0.66 | 0.12 | -33.79 | **< 0.001** |
|  |  | DYS | -0.60 | 0.24 | -14.90 | **< 0.001** |
|  | Sup motor area | CON | -0.55 | 0.11 | -30.17 | **< 0.001** |
|  |  | DYS | -0.47 | 0.27 | -10.14 | **< 0.001** |
|  | Thalamus | CON | -0.47 | 0.19 | -14.70 | **< 0.001** |
|  |  | DYS | -0.38 | 0.31 | -7.24 | **< 0.001** |
| **Supplementary Table 4.** Mean LIs, standard deviations, and one-sample tests against zero using original values.  These tests confirm that both the dyslexia group and the control group are lateralized for each ROI. | | | | | | |

**E. Independent-samples t-tests comparing laterality indices (LIs) between the dyslexia and control groups using original LI values**

| **Task** | **ROI** | ***t*-value** | **df** | ***P*-value** | | **Effect size (d)** |
| --- | --- | --- | --- | --- | --- | --- |
| **Reading task** | Composite ROI | -1.75 | 50.64 | **0.043** | | -0.42 |
|  | Fusiform gyrus | -3.81 | 43.29 | **0.001** | | -0.91 |
|  | Inf frontal gyrus | -1.93 | 56.51 | **0.030** | | -0.46 |
|  | Inf occipital gyrus | -1.87 | 55.92 | **0.033** | | -0.45 |
|  | Inf parietal gyrus | -1.97 | 63.85 | **0.027** | | -0.47 |
|  | Mid temporal gyrus | -1.08 | 64.10 | 0.143 | | -0.26 |
|  | Precentral gyrus | -0.55 | 62.53 | 0.291 | | -0.13 |
| **Rhyming task** | Composite ROI | -1.56 | 39.87 | 0.064 | | -0.37 |
|  | Cerebellum | 0.94 | 65.49 | 0.176 | | 0.22 |
|  | Inf frontal gyrus | -1.53 | 40.37 | 0.067 | | -0.37 |
|  | Mid temporal gyrus | -1.11 | 54.13 | 0.135 | | -0.27 |
|  | Precentral gyrus | -1.65 | 37.62 | 0.054 | | -0.39 |
|  | Sup motor area | -1.82 | 56.59 | **0.037** | | -0.43 |
| **Fluency task** | Composite ROI | -1.31 | 37.67 | 0.100 | | -0.31 |
|  | Cerebellum | 0.96 | 53.90 | 0.170 | | 0.23 |
|  | Inf frontal gyrus | -1.26 | 38.41 | 1.107 | | -0.30 |
|  | Precentral gyrus | -1.33 | 49.13 | 0.096 | | -0.32 |
|  | Sup motor area | -1.64 | 44.32 | 0.054 | | -0.39 |
|  | Thalamus | -1.47 | 54.72 | 0.074 | | -0.36 |

**Supplementary table 5.** Statistics from t-tests comparing the LI values for the dyslexia and control group using original LI values (see Supplementary table 1). Note that the dyslexia group appear less lateralized for several regions, when not controlling for direction of asymmetry, most notably in the reading task.
